# RSV protects bystander cells against IAV infection by triggering secretion of type I and type III interferons

**DOI:** 10.1101/2021.10.11.463877

**Authors:** Maciej Czerkies, Marek Kochańczyk, Zbigniew Korwek, Wiktor Prus, Tomasz Lipniacki

## Abstract

We observed the interference between two prevalent respiratory viruses, respiratory syncytial virus (RSV) and influenza A virus (IAV, H1N1), and characterized its molecular underpinnings in alveolar epithelial cells (A549). We found that RSV induces higher interferon (IFN) β production than IAV and that IFNβ priming confers higher protection against infection with IAV than with RSV. Consequently, we focused on the sequential infection scheme: RSV-then-IAV. Using the A549 WT, IFNAR1 KO, IFNLR1 KO, and IFNAR1–IFNLR1 double KO cell lines we found that both IFNβ and IFNλ are necessary for maximum protection against subsequent infection. Immunostaining revealed that preinfection with RSV partitions the cell population into a subpopulation susceptible to subsequent infection with IAV and an IAV-proof subpopulation. Strikingly, the susceptible cells turned out to be those already compromised and efficiently expressing RSV, whereas the bystander, interferon-primed cells are resistant to IAV infection. Thus, the virus–virus exclusion at the cell population level is not realized through a direct competition for a shared ecological niche (single cell) but rather achieved with the involvement of specific cytokines induced within the host innate immune response.

**Importance:** The influenza A virus (IAV) and the respiratory syncytial virus (RSV) are common recurrent respiratory infectants, which show a relatively high coincidence. We demonstrated that preinfection with RSV partitions the cell population into a subpopulation susceptible to subsequent infection with IAV and an IAV-proof subpopulation. The susceptible cells are those already compromised and efficiently expressing RSV, whereas the bystander cells are resistant to IAV infection. The cross-protective effect critically depends on IFNβ and IFNλ signaling and thus ensues when the proportion of cells preinfected with RSV is relatively low yet sufficient to trigger a pervasive antiviral state in bystander cells. Our study suggests that mild, but not severe, respiratory infections may have a short-lasting protective role against more dangerous respiratory viruses, including SARS-CoV-2.

## Introduction

Human respiratory tract infections are predominantly caused by viruses. Out of a plethora of respiratory viruses, recently expanded by the novel beta coronavirus SARS-CoV-2 (1), the three most common recurrent infectants, originating from different viral families, are rhinovirus (human RV, HRV), influenza A virus (IAV), and the respiratory syncytial virus (RSV, human orthopneumovirus) (2), with the latest being the most common pathogen in severe disease in young children and the elderly (3–5). In temperate areas of the northern hemisphere, the incidence of respiratory infections surges in the winter season and subsides during summer. Detailed examination of temporal patterns of disease outbreaks reveals not only seasonal co-incidence but also apparent avoidance of some types of cocirculating respiratory viruses (6–9). It is unclear whether the epidemiological observations are population-level consequences of an interplay of multiple viral cooperation and competition mechanisms that operate at a single-host level (10). Intriguingly, the epidemiological avoidance patterns are visible also for taxonomically different groups of viruses, suggesting their potential reliance on mechanisms that are not related to specific antibody-mediated cross-protection. Although viral interference can be readily observed *in vitro* (11), its underlying mechanisms are necessarily inextricably intertwined with the response of the host organism and await elucidation from a systemic perspective.

Non-antigenic competitive interactions among respiratory viruses have been suggested based on clinical data analysis (12, 13) and were recapitulated within animal models (14, 15). Two very recent studies (16, 17) demonstrated that an innate immunity mechanism, specifically antiviral cytokine signaling, is implicated in the interference between RV and IAV: blocking the type I interferon (IFN) response restores IAV replication following RV infection. Whereas type I IFNs (IFNα and IFNβ) were originally discovered specifically because of their activity against the influenza virus (18), their roles in RSV infection and disease have been controversial (19–25). A recent animal model study (14) suggested that the RSV–IAV viral interference is antigen-independent, however direct involvement of IFNs has not been demonstrated.

Type I IFNs, together with the more recently identified type III IFNs (IFNλ1–λ4) (26, 27), are produced and secreted by infected cells to trigger transcription of an array of IFN-stimulated genes (ISGs) in both the infected and the bystander cells (28–30). Elevated ISGs restrict viral replication until virus-specific host defense mechanisms develop (31). IFNα and IFNβ additionally attract specific immune system cells to the site of inflammation (22, 32). IFNλ acts more locally, preferentially at epithelial barriers (33), and differentially affects tissue homeostasis: while IFNλ confers longer lasting protection than IFNα (34), chronic IFNλ exposure inhibits proliferation of respiratory epithelial cells, impairing post-infection lung regeneration (35, 36). A significant relative potency of IFNλ in induction of an antiviral state in response to IAV has been demonstrated (34, 37–40). In the absence of IFNβ, IFNλ has been proposed to be the primary factor preventing RSV infection (25).

Propagation of RSV strongly depends on the expression of its two nonstructural proteins, NS1 and NS2, that, among multiple modalities of interference with the innate immune response, block the synthesis of interferons and downregulate the expression of ISGs (by inhibition of STAT1/2 signaling) (41, 42). Of note, the strength of the immune response to RSV increases with the contents of the immunostimulatory defective viral genomes (DVGs), typically incomplete viral progeny that can stimulate production of interferons (43, 44). DVGs are naturally generated during RSV replication and may limit the spread of RSV as well as other coinfecting viruses. *In vitro*, at low multiplicities of infection, when the selective pressure on virus is high, the incidence of DVGs is low (43). The kinetics of DVGs accumulation may predict the clinical outcome of an RSV infection in humans; detection of DVGs early after infection has been associated with low viral loads and mild disease (45).

In this study, we sought to determine molecular underpinnings of the interactions between RSV and IAV. We used a human alveolar epithelial cell line (A549) and characterized direct inter-viral relations and interferon-mediated interference *in vitro* at the single-cell level. We found that RSV induces higher IFNβ production than IAV and that IFNβ is a more potent inducer of STAT1/2 signaling than IFNλ1. Moreover, IFNβ-stimulated cells are more resistant to infection with IAV than to infection with RSV. In light of these findings, we focused on the sequential coinfection scheme, in which cells are preinfected with RSV and then infected with IAV. We developed A549-derived cell lines with knockout (KO) of type I IFN receptor (IFNAR1 KO), type III IFN receptor subunit (IFNLR1 KO), and double KO (IFNAR1–IFNLR1 dKO). They enabled us to find that IFNβ is the main inducer of an antiviral state in an RSV-infected population of A549 cells and both interferons, IFNβ and IFNλ, are simultaneously necessary for building maximum protection against a subsequent infection with IAV. Immunostaining revealed that preinfection with RSV partitions the cell population into a subpopulation susceptible to subsequent infection with IAV and an IAV-proof subpopulation. Strikingly, the susceptible cells turned out to be those already compromised and efficiently expressing the priming virus. Overall, we conclude that in the sequential RSV-then-IAV infection scheme, the virus–virus exclusion at the cell-population level is not realized through a direct competition for a shared ecological niche (single cell) but rather achieved with the involvement of specific cytokines induced within the host innate immune response.

## Results

### IFNβ and IFNλ mediate STAT1/2 activation upon RSV and IAV infection

We first characterized induction of the antiviral state by interferon signaling. We found that RSV is a more potent inducer of IFNβ than IAV at comparable MOIs during 48 hours post-infection (*cf*. Fig. 1A, Fig. 1B). As shown in Fig. 1C, upon infection with RSV at low MOI, only a tiny fraction of cells (1–5%) produces IFNβ, which is however sufficient to activate the STAT pathway in remaining cells. Cells expressing IFNβ typically do not express RSV proteins which may result from the early stage of infection, suppression of RSV protein expression by innate immune response in infected cells, or infection by DVGs. At the same time, in cells that express RSV proteins, non-structural proteins may inhibit synthesis of IFNβ at various levels (46–50). Detection of viral RNA within an RSV-infected cell is manifested by activation of IRF3 (which gets phosphorylated and then translocates to the nucleus becoming visible in images from immunostaining). Propagation of RSV and IAV for 48 hours from MOI = 0.01, induction of STAT1 activity and its inhibition over time are illustrated in Supplementary Fig. S1 and Fig. S2, respectively.

**Figure 1.**
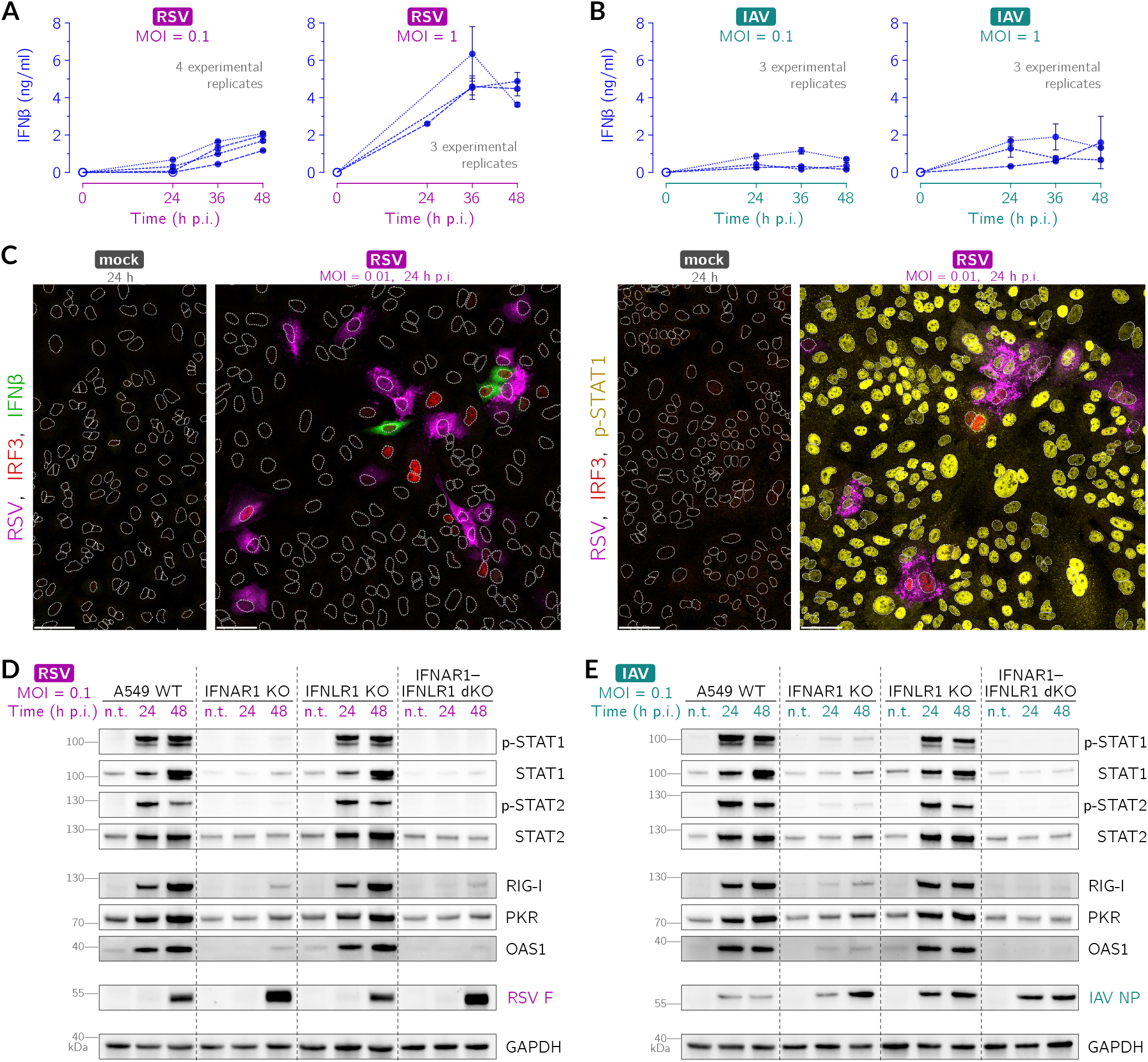
IFNβ and IFNλ mediate STAT1 and STAT2 activation upon RSV and IAV infection in A549 cells. (**A, B**) ELISA for IFNβ following (A) RSV and (B) IAV infection, both at MOI = 0.1 and MOI = 1. Open circles mark data points with cytokine concentration below detection range. Error bars are s.d. of technical replicates. **(C)** IFNβ (left subpanels, green) and activation of STAT1 (right subpanels, yellow) and IRF3 (both subpanels, red) 24 hours after infection of A549 WT cells with RSV (magenta, polyclonal anti-RSV antibody) at MOI = 0.01. White dotted lines are nuclear outlines determined based on DAPI counterstaining (channel not shown). Scale bar, 50 μm. (**D, E**) Activation of STAT1, STAT2, and interferon-stimulated genes (RIG-I, PKR, OAS1) upon (C) RSV and (D) IAV infection, both at MOI 0.1, at 24 and 48 hours post-infection (h p.i.), in A549 WT, IFNAR1 KO, IFNLR1 KO, and IFNAR1–IFNLR1 dKO cells. Non-treated cells are labeled n.t. Activated STATs, p-STAT1 and p-STAT2, are STAT1 and STAT2 phosphorylated at Tyr 701 and at Tyr 690, respectively. RSV F, RSV fusion glycoprotein; IAV NP, IAV nucleoprotein.

We then analyzed interferon-mediated activation of STAT1/2 during RSV or IAV infection and consecutive induction of ISGs that are the primary correlates of the antiviral state: RIG-I (viral RNA sensor (51, 52)), PKR (indirect translation inhibitor (53)), and OAS1 (direct activator of RNase L, the RNA eradicator (54)). We juxtaposed results for A549 WT cells with those for the A549-derived cell lines in which we knocked out IFNα/β receptor IFNAR1, heterodimeric IFNλ receptor subunit IFNLR1, and both these genes (Fig. 1D, Fig. 1E). We observed that infection with RSV or IAV leads to robust activation (phosphorylation) of STAT1 and STAT2 in A549 WT cells as well as in IFNLR1 KO cells. In IFNAR1–IFNLR1 dKO cells, there is no observable STAT1/2 phosphorylation, whereas in IFNAR1 KO cells, secreted type III IFNs (IFNλ) leads to relatively weak STAT1/2 phosphorylation. The observed patterns of STAT activity translate directly to the patterns of accumulation of RIG-I, PKR, and OAS1, as well as STAT1 and STAT2, with the exception of IFNAR1–IFNLR1 dKO cells, in which after RSV we observe an increase of RIG-I and PKR, which is apparently independent of STAT1–STAT2 signaling. Cells devoid of the receptor for IFNβ, but not cells devoid of the receptor for IFNλ, show higher levels of RSV fusion glycoprotein, suggesting faster virus replication (Fig. 1D). Of note, the lack of either IFNβ receptor or IFNλ receptor facilitates IAV replication, despite IFNLR1 KO having minimal impact on STAT1/2 signaling and accumulation of ISGs (Fig. 1E). Altogether, we conclude that despite divergent efficacy of IFNβ induction by RSV and IAV, upon infection with any of these viruses, IFNβ is the main activator of STAT1 and STAT2 signaling. We did not analyze the impact of IFNγ, because the lack of STAT1 activation in IFNAR1 KO cells and in IFNAR1–IFNLR1 dKO cells (Fig. 1D, Fig. 1E) suggests that this cytokine is not released by the studied cells and as such does not contribute to the inhibition of viral proliferation in our experimental setup.

### Prestimulation with IFNβ inhibits propagation of RSV and IAV

To estimate the impact of priming with IFNβ and IFNλ1 on the infectiousness and propagation of RSV and IAV, we quantified viral RNA molecules per cell and analyzed the abundance of viral proteins. We performed infection experiments without or with a preceding, day-long stimulation with IFNβ (1000 U/ml) or with IFNλ1 (50 ng/ml). In Fig. 2, we juxtaposed results for these two stimulation protocols. We found that 24 hours post-infection at MOI 0.1, prestimulation with IFNβ limits RSV progeny by nearly one order of magnitude, whereas IAV progeny is reduced by at least two orders of magnitude (*cf*. Fig. 2A, Fig. 2B). At MOI 1, when nearly all cells are infected, prestimulation with IFNβ limits RSV proliferation about five-fold, while the relative reduction of IAV proliferation is as significant as in the case of the lower MOI (*cf*. Fig. 2A, Fig. 2B). Prestimulation with IFNβ triggers the antiviral state manifested by accumulation of ISGs: RIG-I, PKR, and OAS1 and STAT1/2. These proteins maintain their elevated levels 48 hours post-infection (Fig. 2C, Fig. 2D). Consequently, viral proteins are less abundant in the IFNβ-prestimulated cells than in the non-prestimulated cells. We conclude results shown in Fig. 1 and Fig. 2 by stating that both IFNβ and IFNλ secreted by infected cells are responsible for confinement of the spread of IAV, however only IFNβ appears to be capable of attenuating the propagation of RSV.

**Figure 2.**
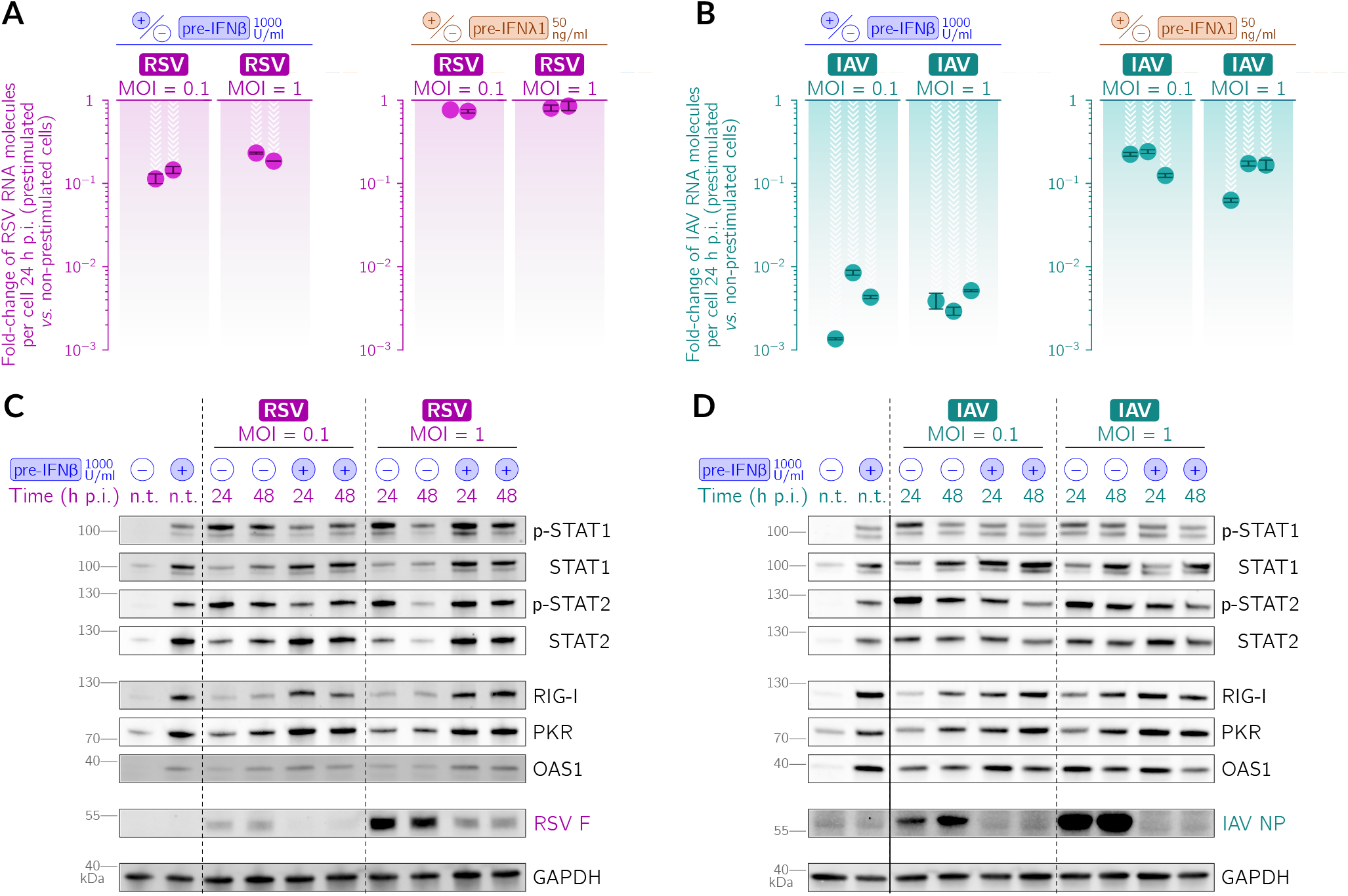
Prestimulation with IFNβ or IFNλ1 inhibits propagation of RSV and IAV in A549 WT cells. (**A, B**) Effect of prestimulation with IFNβ (1000 U/ml) or IFNλ1 (50 ng/ml) 24 hours prior to infection on the relative abundance of viral RNA 24 hours post (A) RSV and (B) IAV infection, both at MOI = 0.1 and MOI = 1. Each filled circle corresponds to fold-change w.r. to non-prestimulated cells in a single experimental replicate (note the logarithmic scale). Error bars show the smallest and the largest ratio of the non-prestimulated to IFN-prestimulated cells for all pairs of respective technical replicates. (**C, D**) Effect of prestimulation with IFNβ (1000 U/ml) 24 h prior to infection on the abundance and activation of STAT1 and STAT2 and abundance of interferon-stimulated genes (RIG-I, PKR, OAS1) and (C) RSV fusion glycoprotein (RSV F) and (D) IAV nucleoprotein (IAV NP), 24 and 48 hours post-infection (h p.i.), both at MOI = 0.1 and MOI = 1. Non-treated cells are labeled n.t. Activated STATs, p-STAT1 and p-STAT2, are STAT1 and STAT2 phosphorylated at Tyr 701 and at Tyr 690, respectively.

### Impact of RSV or IAV infection on STAT1 activation

To analyze at the single-cell level how infected cells induce the antiviral state in bystander cells, we performed infections at MOI 0.1, fixed the cells 24 hours post-infection, and immunostained them for active IRF3 (as an evidence of detection of viral RNA by cell), p-STAT1 (used as an indication of the antiviral state), and viral proteins. In two-channel overlays shown in Fig. 3A, Fig. 3B, it can be noticed that p-STAT1 is present in most bystander cells (at various levels), whereas the infected cells exhibit p-STAT1 only sporadically. This effect may indicate the capability of the viruses to suppress the response to IFN stimulation (46–50). We quantified this effect by looking directly at the presence of viral proteins as the definite manifestation of productive infection, and at nuclear p-STAT1 as an indication of antiviral alertness. Based on the expression of RSV proteins or IAV nucleoprotein, we classified cells as either virus-positive or virus-negative and collected histograms of nuclear p-STAT1 intensity in each cell within the two groups. In Fig. 3C, Fig. 3D we can see that the two distributions are indeed different for virus-positive and virus-negative cells, with on average higher STAT1 phosphorylation in virus-negative cells. This indicates that the antiviral state is more pronounced in bystander uninfected cells than in RSV-infected or IAV-infected cells 24 hours post-infection. To express the extent to which the two distributions are disjoint, we calculated the Kolmogorov–Smirnov (KS) statistic. KS attains 1 for fully disjoint distributions and 0 for exactly overlapping distributions. Across all experimental replicates, KS lies in the range 0.42–0.58 (95% CrI: 0.39–0.67) in the case of infection with RSV and in the range 0.57–0.59 (95% CrI: 0.51–0.68) in the case of infection with IAV. See also Supplementary Fig. S1 and Fig. S2 documenting a decline of STAT1 phosphorylation in viral proteins-expressing cells.

**Figure 3.**
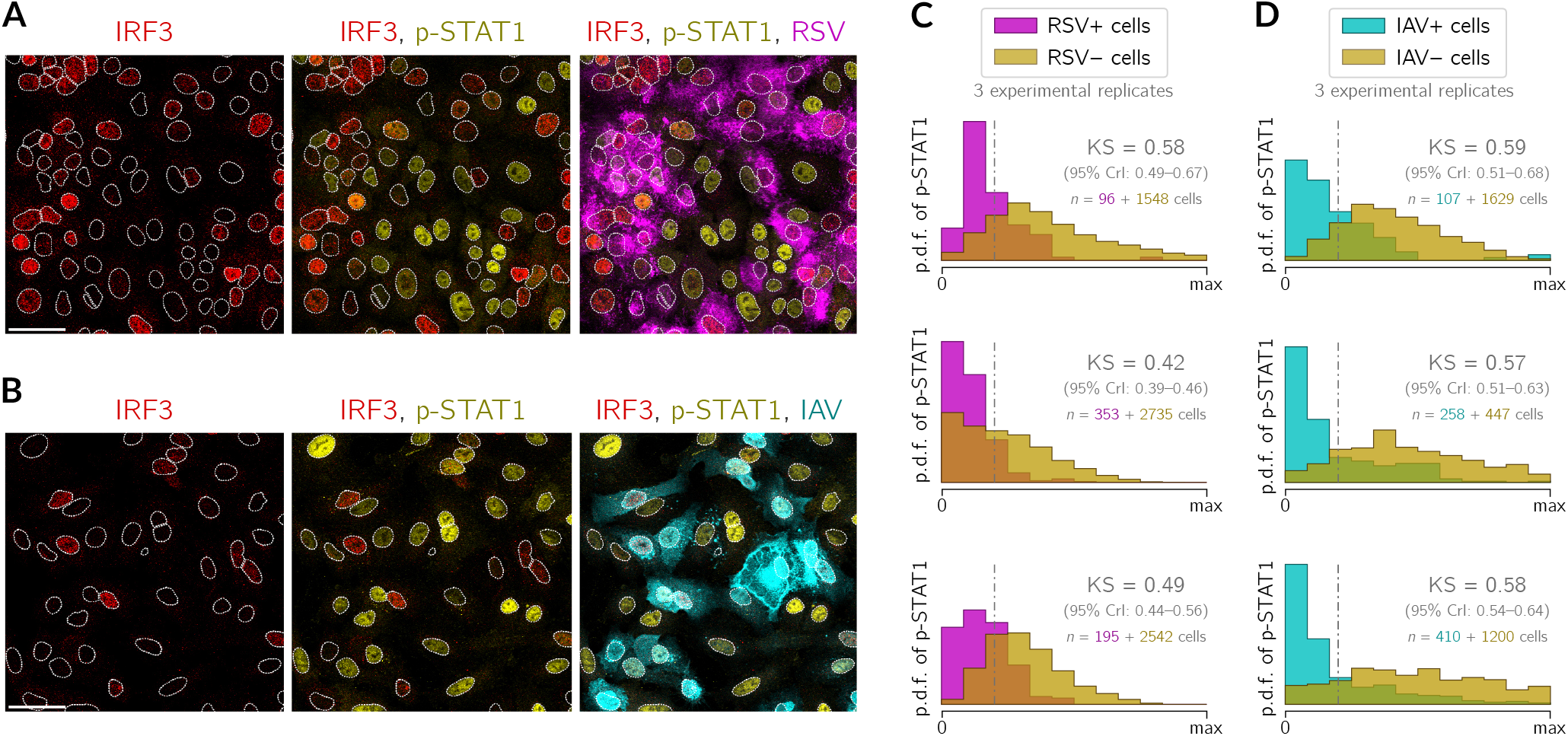
Impact of RSV or IAV infection on STAT1 activation. (**A, B**) A549 WT cells 24 hours post-infection with (A) RSV at MOI = 0.1 or (B) IAV at MOI = 0.1. Channels with IRF3 (red), p-STAT1 (yellow), RSV (magenta, polyclonal anti-RSV antibody, only in panel A) or IAV (cyan, anti-IAV nucleoprotein antibody, only in panel B) are shown as incremental overlays. White dotted lines are nuclear outlines determined based on DAPI counterstaining (channel not shown). Scale bar, 50 μm. (**C, D**) Histograms showing the empirical probability density function (p.d.f.) of p-STAT1 24 hours postinfection with (C) RSV at MOI = 0.1 or (D) IAV at MOI = 0.1 in cells classified as either expressing (+) or not expressing (−) the virus. In each pair of overlaid histograms, a Kolmogorov–Smirnov statistic is reported (with its credible interval, CrI) and a corresponding maximally discriminating threshold is marked with a vertical dash-dotted line. In each subpanel of (C) and (D), three experimental replicates are shown separately and the numbers of cells that were classified as either virus-positive or virus-negative are given on the left- and the right-hand side of the plus sign, respectively.

### Effect of prestimulation with IFNβ or IFNλ1 on IAV spread in A549 WT cells

Thus far, we have used cell-population techniques to show that IFNβ is a more potent inducer of the antiviral state than IFNλ1 (Fig. 1) and, as a consequence, in A549 cells viral proliferation is more suppressed after prestimulation with IFNβ than with IFNλ1 (Fig. 2A, 2B). Since RSV turned out to elicit stronger IFNβ response than IAV, to characterize the potential interferon-mediated cross-protection between the two viruses we decided to investigate specifically the dual infection scheme in which the cells are first preinfected with RSV and after 24 hours infected with IAV.

To gain insight into the interferon-mediated interaction at the single-cell level, we quantified the protective effect of IFNβ or IFNλ1 in the context of IAV infection based on imaging data (Fig. 4A). In Fig. 4B we show proportions of cells expressing viral nucleoprotein 24 hours after treatment with IAV at MOI = 1 in each field of view in the case of non-prestimulated A549 WT cells and A549 WT cells that were prestimulated with either IFNβ or IFNλ1. Based on image quantifications for two experimental replicates, we found that 40–45% of non-prestimulated cells express nucleoprotein 24 hours after infection. This proportion drops to about 3% after prestimulation with IFNβ and to about 10% after prestimulation with IFNλ1. Relations between these proportions provide additional evidence for strong attenuation of the IAV spread conferred by IFNs in A549 cells.

**Figure 4.**
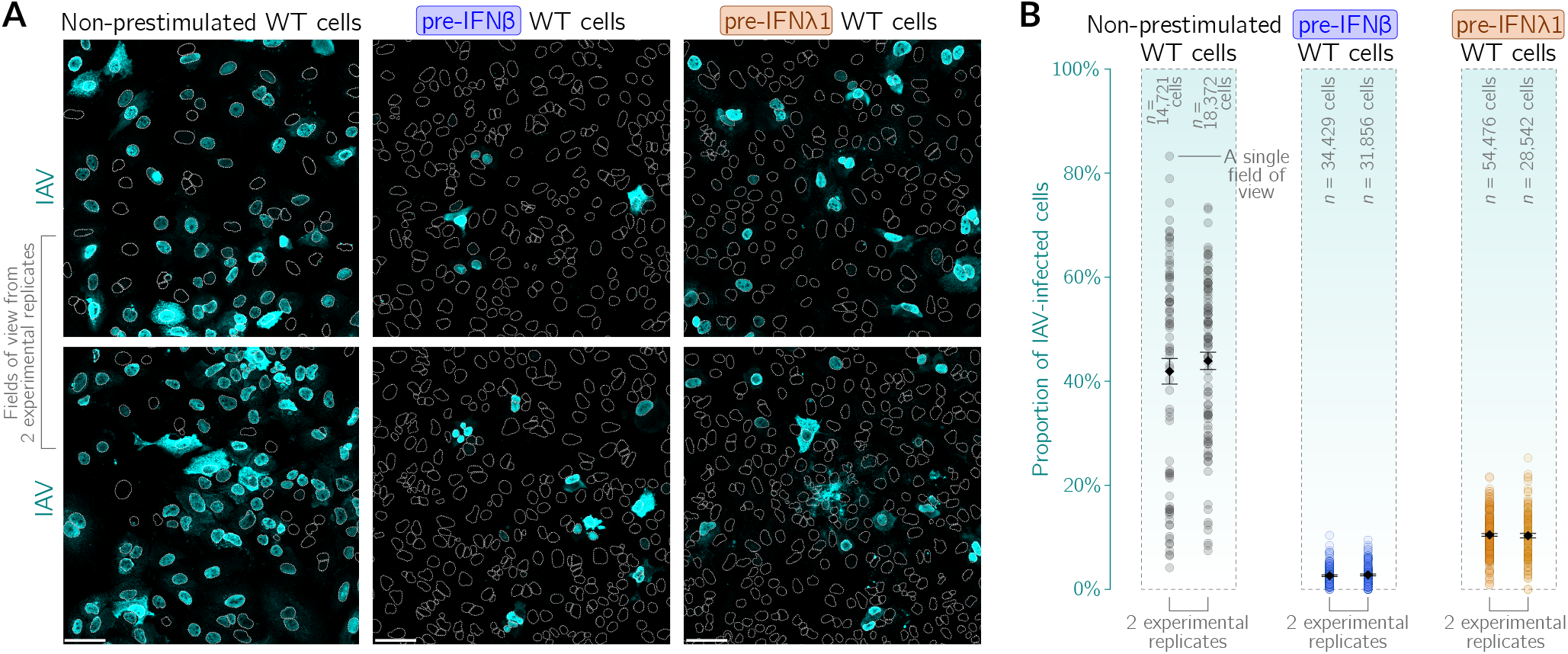
Effect of prestimulation with IFNβ or IFNλ1 on IAV spread in A549 WT cells. Non-prestimulated A549 WT cells or A549 WT cells prestimulated with either IFNβ (100 U/ml) or IFNλ1 (50 ng/ml) for 24 hours were infected with IAV at MOI = 1 and fixed 24 hours post-infection (i.e., 48 hours since the beginning of prestimulation). Interferon-containing cell culture media were not displaced upon infection. **(A)** Representative fields of view showing non-prestimulated cells and cells prestimulated with either IFNβ or IFNλ1 after infection with IAV (cyan, anti-IAV nucleoprotein antibody). White dotted lines are nuclear outlines determined based on DAPI counterstaining (channel not shown). For each pretreatment condition, sample images from two experimental replicates are shown. Scale bar, 50 μm. **(B)** Proportions of productively IAV-infected cells among non-prestimulated cells and cells prestimulated with either IFNβ or IFNλ1, based on nucleoprotein expression. Data from two independent experiments are shown separately; each circle corresponds to a single field of view. The fields of view do not overlap. Black diamonds and featuring error bars denote mean proportions ± s.e.m. The number of cells analyzed in all fields of view from each experiment is given above respective data series.

### RSV preinfection inhibits IAV infection in an interferon-dependent manner

In the course of viral infection there are cells that are initially infected by the primary virus and bystander cells that, in response to IFN secreted by the initially infected cells, may develop the antiviral state and become partially protected from secondary infections. Our prior experiments with IFN prestimulation and high IAV MOI allowed us to characterize the response of the cells that had uniformly developed the antiviral state and were then challenged with the virus, which mimics only one subpopulation of the cells that appears during ongoing infection. Since in our dual infection scheme we wanted to apply the second virus to a cell population that consist of the subpopulation of the preinfected and IFN-producing cells and the subpopulation of not (yet) infected but IFN-stimulated cells, we primed cells with the first virus, RSV, at MOI = 0.1 and after 30 hours applied the second virus, IAV, at MOI = 1. To corroborate and discern the relative importance of IFNβ or IFNλ in their native RSV-elicited mixture, we used additionally A549 IFNAR1 KO, IFNLR1 KO, and IFNAR1–IFNLR1 dKO cell lines.

In Fig. 5A and Fig. 5B, we show imaging-based single-cell level analysis of the cell population after dual infection. After preinfection with RSV, the proportion of WT A549 cells infected with the second virus, IAV, is reduced by an order of magnitude with respect to non-preinfected cells (*cf*. non-prestimulated WT cells in Fig. 4B and RSV-primed WT cells in Fig. 5B, which gives a tenfold reduction of IAV nucleoprotein-expressing cells, from ~40% to ~4%). The protective effect conferred by RSV-priming is significantly weaker in IFNAR1 KO cells incapable of responding to IFNβ (reduction from ~40% to ~20%) and only slightly weaker in IFNLR1 KO cells incapable of responding to IFNλ (reduction from ~40% to ~10%). The priming effect is virtually non-existent in cells devoid of receptors for both IFNβ and IFNλ. According to dPCR measurements shown in Fig. 5C, the amount of IAV RNA in a population of RSV-preinfected WT cells is 10–20 times lower than in a non-preinfected population of WT cells. Additionally, in populations of RSV-preinfected cells from three KO cell lines, the amount of IAV RNA is higher than in the WT cell population, and the ratios of IAV RNA expression between these lines are similar to the ratios of IAV-infected cell proportions shown in Fig. 5B. Since IFNs block IAV at the level of replication of viral RNA and synthesis of viral proteins, the accumulated effect at the level of viral proteins may be expected to be more pronounced.

**Figure 5.**
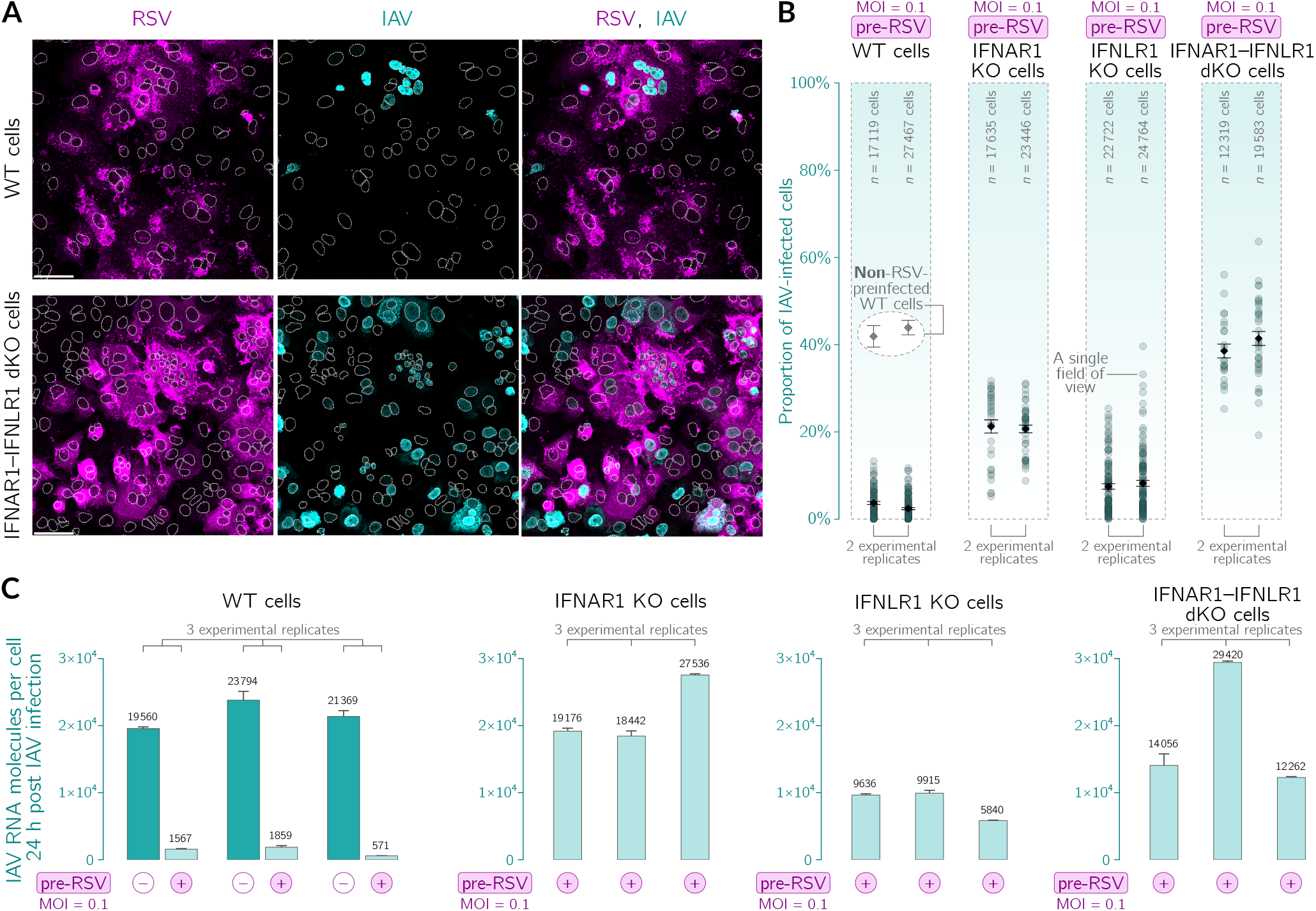
RSV preinfection inhibits IAV infection in an interferon-dependent manner in A549 cells. Analyzed are infections in four A549 cell lines: WT, IFNAR1 KO, IFNLR1 KO, and IFNAR1–IFNLR1 double KO (dKO). Non-preinfected cells and cells preinfected with RSV at MOI = 0.1 for 30 hours were infected with IAV at MOI = 1 and cultured subsequently for 24 hours. Cell culture media was not displaced upon IAV infection. **(A)** Representative fields of view showing WT and IFNAR1–IFNLR1 dKO cells preinfected with RSV (magenta, polyclonal anti-RSV antibody) after infection with IAV (cyan, anti-IAV nucleoprotein antibody). White dotted lines are nuclear outlines determined based on DAPI counterstaining (channel not shown). Scale bar, 50 μm. **(B)** Proportions of IAV-infected cells in an RSV-preinfected cell population. Data from two independent experiments are shown separately; each circle corresponds to a single field of view. Black diamonds and featuring error bars denote mean proportions ± s.e.m. Two encircled gray diamonds above the data points for WT cells are mean proportions of the IAV-infected WT cells that were not subjected to preinfection with RSV (non-prestimulated WT cells in Fig. 4B), shown here again, as a reference. **(C)** Impact of preinfection with RSV on the number of IAV RNA molecules per cell in all four considered cell lines. Cells were infected with RSV and, after 30 hours, with IAV. dPCR has been performed for cells collected 24 hours post IAV (i.e., 54 hours post RSV) infection. Error bars are s.d. of technical replicates.

In Fig. 6A and Fig. 6B, we demonstrate that the protective effect conferred by RSV-priming depends on RSV MOI. The strongest protection is achieved for intermediate RSV MOI of ~0.1– 0.3, whereas for RSV MOI of ~1, protection is a bit weaker and for RSV MOI of ~0.01 it is much weaker (Fig. 6B). At the lowest considered RSV MOI, a small number of cells secrete IFNs, which is likely insufficient to evoke the pervasive antiviral state in bystander cells. We note that varying pre-RSV MOI is somewhat similar to changing the time interval between subsequent infections, because longer intervals permit more secondary infections with the priming virus and higher accumulation of interferons.

**Figure 6.**
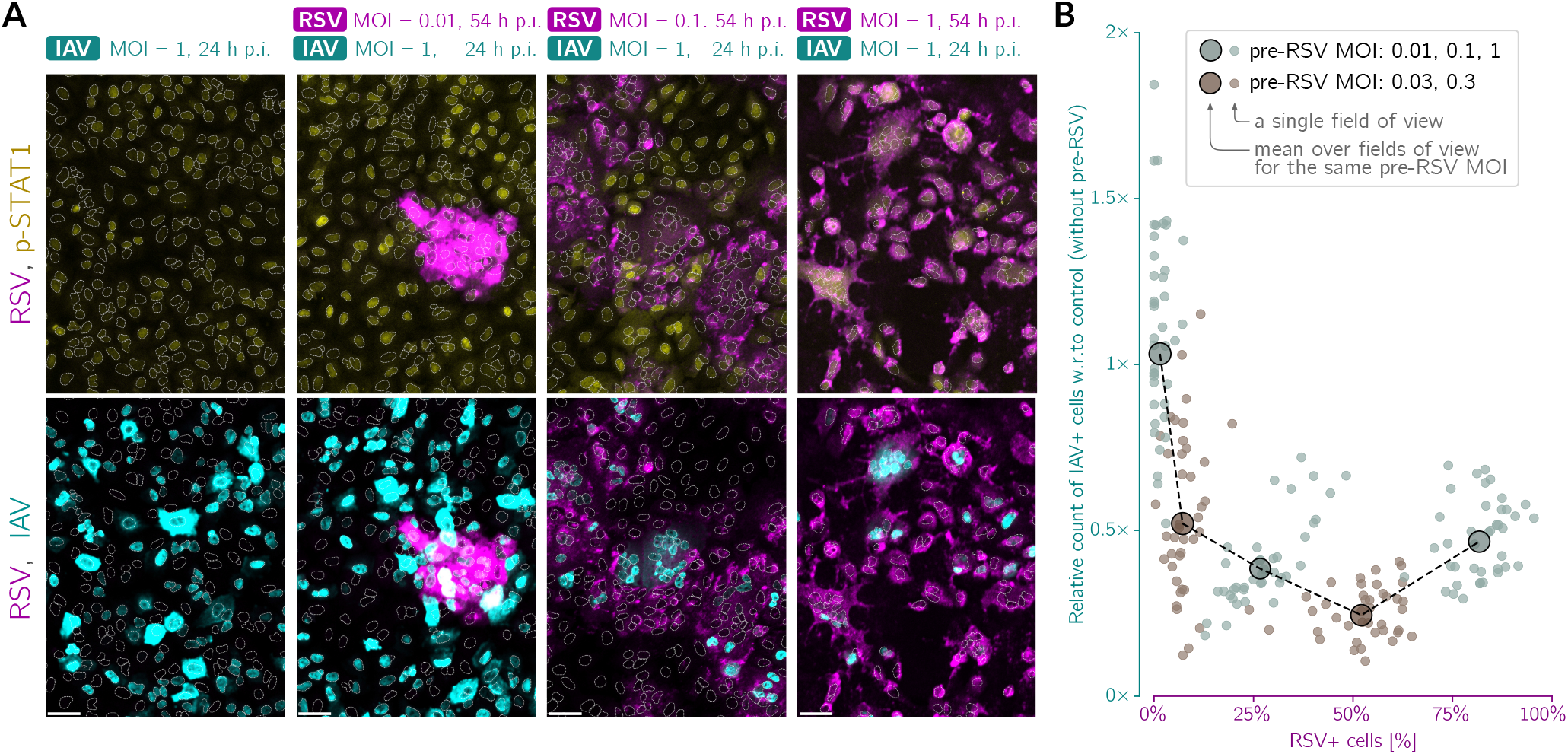
RSV preinfection inhibits IAV infection in an RSV MOI-dependent manner. **(A)** Representative fields of view showing A549 WT cells preinfected with RSV at different MOIs (0.01, 0.03, 0.1, 0.3, 1) for 30 hours and then infected with IAV for additional 24 hours. Top row, overlays of RSV (magenta, polyclonal anti-RSV antibody) and p-STAT1 (yellow); bottom row, overlays of RSV (magenta, polyclonal anti-RSV antibody) and IAV (cyan, anti-IAV nucleoprotein antibody). White dotted lines are nuclear outlines determined based on DAPI counterstaining (channel not shown). Scale bar, 50 μm. **(B)** Proportion of the count of IAV-infected cells after preinfection with RSV to the count of IAV-infected cells without preinfection, as a function of the percentage of cells expressing RSV proteins. Small circles correspond to individual fields of view, whereas large circles represent values averaged over all fields of view for a given MOI of applied pre-RSV. Data from two representative experiments; the RSV stock used in these experiments (performed in revision) was different than in experiments shown in Fig. 1, Fig. 2, Fig. 3, Fig. 5, and Fig. 7.

Overall, this demonstrates that preinfection with RSV protects from infection with IAV in an interferon-dependent manner in A549 cells. The protective effect may be attributed nearly entirely to the paracrine stimulation with both IFNβ and IFNλ, with a major contribution from IFNβ.

### IAV infects preferentially the cells that express RSV proteins

Visual inspection of WT cells suggested that IAV nucleoprotein is preferentially expressed in the cells with apparent expression of RSV proteins (see Fig. 7A). To quantitatively verify this hypothesis, as in Fig. 3D, we classified cells as virus-positive and virus-negative. We decided to use the measured level of IAV NP as the binary criterion because of its mostly nuclear location, which facilitates image analysis. We then collected histograms of intensity of RSV proteins in the cells belonging to these two groups. Clearly, in A549 WT cells, the IAV+ cells are associated with higher abundance of RSV proteins than IAV− cells (KS of 0.62 [95% CrI: 0.599– 0.643] indicates good discernibility). This effect is much weaker in IFNAR1–IFNLR1 dKO cells, which cannot respond to IFNβ or IFNλ (KS of 0.15 [95% CrI: 0.137–0.154] indicates that the two distributions are nearly overlapping). This clearly shows that the RSV–IAV exclusion is not realized through a direct competition for a shared ecological niche, a single cell, but rather achieved with the involvement of interferon signaling.

**Figure 7.**
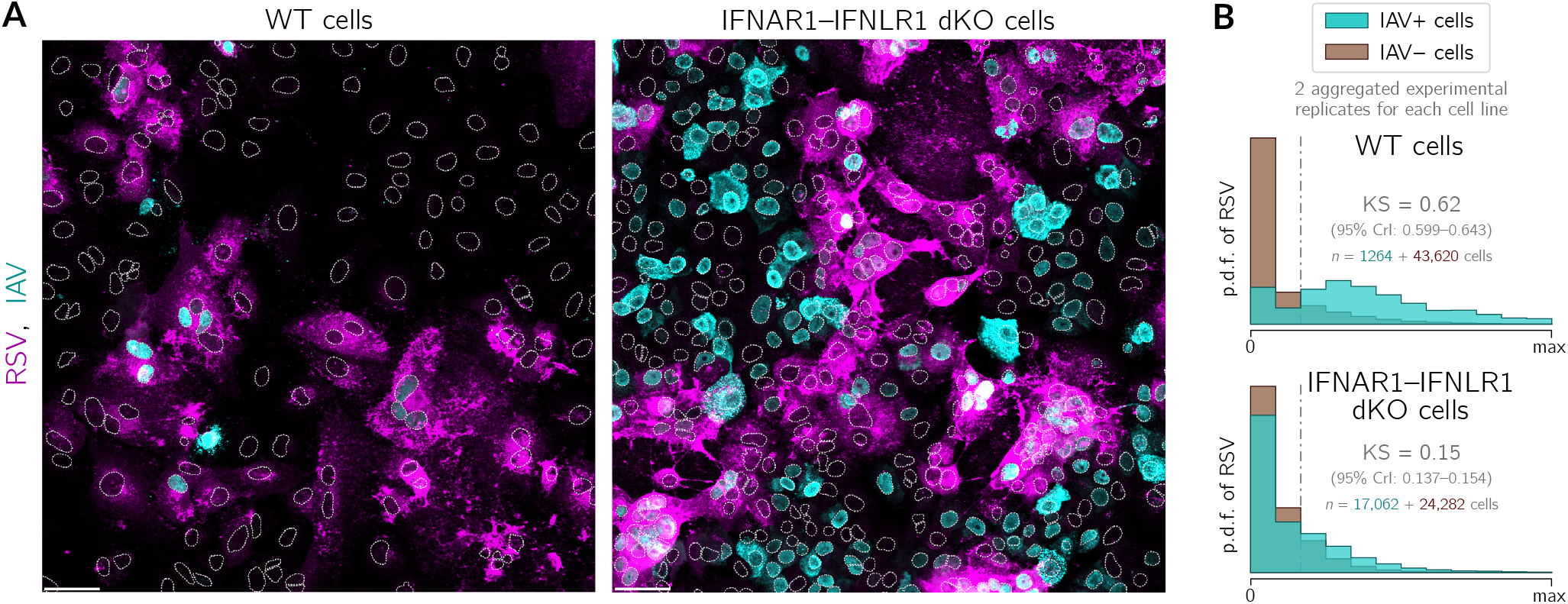
IAV infects preferentially A549 WT cells expressing RSV proteins, but this effect is not observed in A549 IFNAR1–IFNLR1 dKO cells. A549 WT and A549 IFNAR1–IFNLR1 dKO cells were preinfected with RSV (MOI = 0.1) 30 hours before infection with IAV (MOI = 1) and fixed 24 hours after infection with IAV. **(A)** Representative fields of view showing WT cells and IFNAR1–IFNLR1 dKO cells preinfected with RSV (magenta, polyclonal anti-RSV antibody), after subsequent IAV (cyan, anti-IAV nucleoprotein antibody) infection (overlays). White dotted lines are nuclear outlines determined based on DAPI counterstaining (channel not shown). Scale bar, 50 μm. **(B)** Histograms of RSV intensity (p.d.f., probability density function) in cells classified as either infected (cyan) or non-infected (brown) with IAV in A549 WT cells and A549 IFNAR1–IFNLR1 dKO cells. Histograms show aggregated data from two experimental replicates. The numbers of cells that were classified as either virus-positive or virus-negative are given on the left- and the right-hand side of the plus sign, respectively.

Based on distributions shown in Fig. 7B we introduced a threshold at which two distributions are best separable (vertical dot-dashed line), allowing us to stratify cells into RSV-positive and RSV-negative ones. Then we found that in experiments with WT cells, among RSV-positive cells (that constituted 20% of all cells), 11% were IAV-positive, whereas among RSV-negative cells (that constituted 80% of all cells), less than 1% were IAV-positive. In experiments with dKO cells, among RSV-positive dKO cells (that constituted 25% of all cells), 56% were IAV-positive and among RSV-negative cells (that constituted 75% of all cells), as much as 37% were IAV-positive.

To demonstrate that the RSV-and-IAV coinfections observed at the single-cell level are mainly due to IAV infections of cells that already express RSV proteins, we performed an experiment in which 40 hours after infection with RSV (at MOI = 0.1) cells are infected with IAV, for either 10 or 24 hours (Fig. 8). Within this protocol it is less probable that a cell that has not been infected during the first 40 hours becomes infected jointly with two viruses in the last 10 hours of the experiment. Immunostaining images (Fig. 8A, Fig. 8C) clearly show that localization of IAV-infected cells recapitulates the spatial pattern of RSV-infected cell clusters and the cells infected with IAV have a higher chance to be infected with RSV (and *vice versa*). This effect is even stronger 10 hours than 24 hours after infection with IAV (Fig. 8B, Fig. 8D).

**Figure 8.**
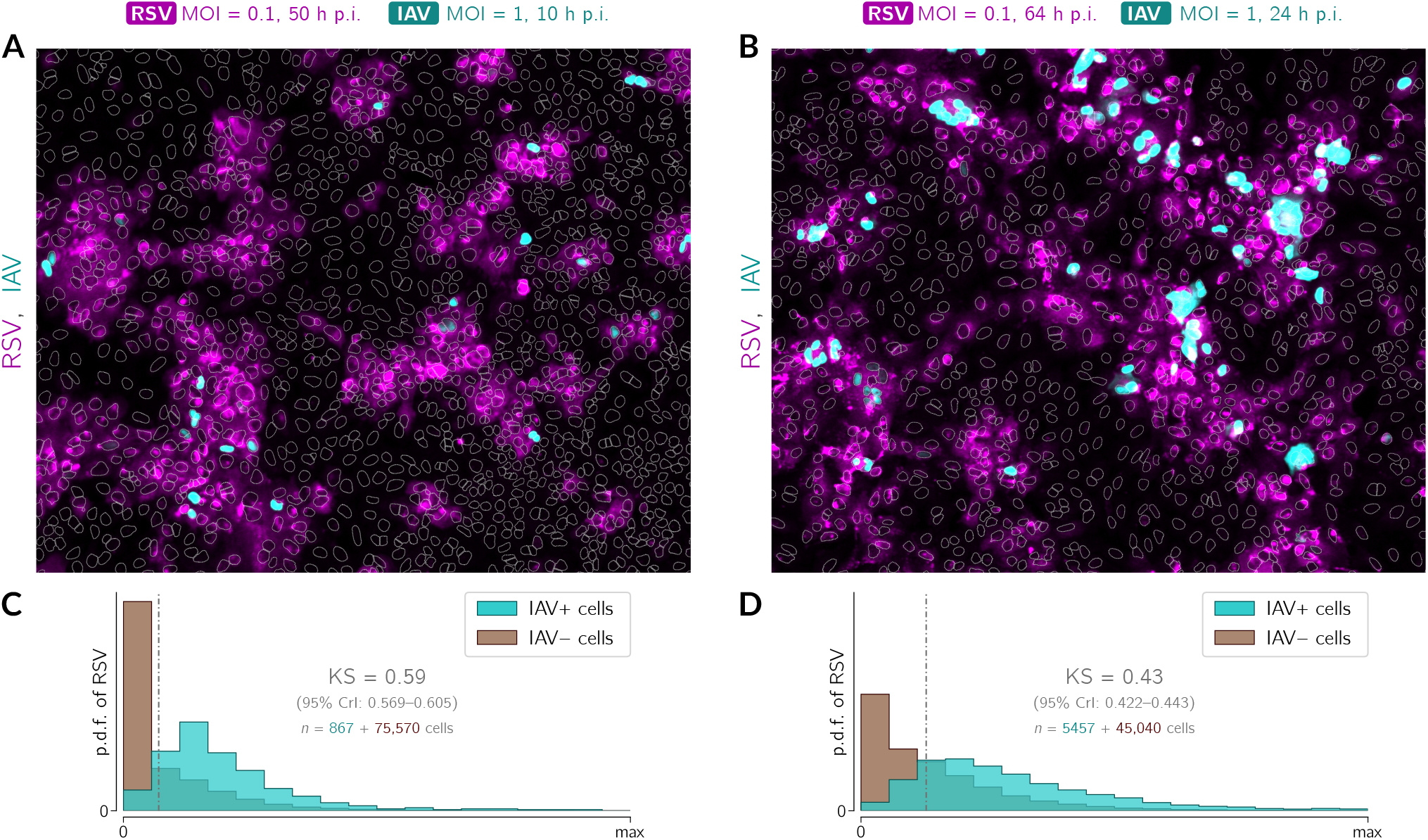
IAV’s preference to infects RSV-preinfected cells is maintained over time. A549 WT cells were preinfected with RSV (MOI = 0.1) 40 hours before infection with IAV (MOI = 1) and fixed 10 or 24 hours after infection with IAV. (**A, B**) Representative fields of view showing cells preinfected with RSV (magenta, polyclonal anti-RSV antibody), after subsequent IAV (cyan, anti-IAV nucleoprotein antibody) infection (overlays) 10 hours (A) and 24 hours (B) after infection with IAV. White dotted lines are nuclear outlines determined based on DAPI counterstaining (channel not shown). Scale bar, 50 μm. (**C, D**) Histograms of RSV intensity (p.d.f., probability density function) in cells classified as either infected (cyan) or non-infected (brown) with IAV. The numbers of cells that were classified as either virus-positive or virus-negative are given on the left- and the right-hand side of the plus sign, respectively. Data from two representative experiments; the RSV stock used in these experiments (performed in revision) was different than in experiments shown in Fig. 1, Fig. 2, Fig. 3, Fig. 5, and Fig. 7.

As in the case of Fig. 7B, we stratified cells into RSV-positive and RSV-negative. Then we found that 10 hours p.i. with IAV (Fig. 8C), among RSV-positive cells (that constituted 34% of all cells), 3% were IAV-positive, whereas among RSV-negative WT cells (that constitute 66% of all cells), about 0.1% were IAV-positive. We also found that 24 hours p.i. with IAV (Fig. 8D), 46% of all cells were RSV-positive and among these cells 20% were IAV-positive; among RSV-negative cells (54%), about 3% were IAV-positive.

## Discussion

To replicate, an invading pathogen has to evade or compromise the innate immunity. When challenged by a virus, the innate immune system acts to impede pathogen replication before a more specific adaptive response is evoked. In the context of sequential viral coinfections, the priming virus both breaks through the innate protection in infected cells and, indirectly, via interferon secreted by these cells, triggers the protective antiviral state in bystander cells. These two antagonistic processes determine the fate of both the primary and the secondary infection at the cell-population level.

To obtain insight into this interplay at the single-cell level, we studied how upon a priming infection with RSV, individual cells become either cross-protected or, conversely, more vulnerable to the second virus, IAV. The choice of these two pathogens has been motivated by the fact that RSV and influenza viruses show relatively high coincidence (55). The particular order of infections that we focused on results from our observations that (1) RSV induces higher IFNβ production than IAV, (2) IFNβ-stimulated cells are more resistant to infection with IAV than to infection with RSV, and (3) IAV is sensitive to IFNλ and this sensitivity does not rely solely on STAT1/2 signaling. We thus inferred that the potential cross-protection may be more pronounced in the RSV-then-IAV protocol, which is consistent with a comparison of RSV-then-IAV and IAV-then-RSV coinfections in a mouse model, showing that the IAV-then-RSV coinfection is associated with higher IAV loads and mouse mortality (56) (see also (57)).

We found that IFNβ is the main inducer of the STAT1/2-associated antiviral state in RSV-infected A549 cells and both interferons, IFNβ and IFNλ, are simultaneously necessary for building maximum protection against a subsequent infection with IAV. A synergistic effect of type I and type III interferon, consistent with our findings, has been previously reported in mice by Mordstein *et al*. (58). In our coinfection experiments, we infected the epithelial alveolar cells with RSV at MOI of 0.1 to readily observe their stratification into the compromised infected cells and the reinforced bystander cells. Immunostaining revealed that preinfection with RSV partitions the cell population into a subpopulation susceptible to a subsequent infection with IAV and an IAV-proof subpopulation. This dynamic functional differentiation of cells may be amplified *in vivo* by differential propensity of specific cell types to permit or restrict viral replication, produce interferons, and build up the antiviral state (59). We found that the susceptible cells turned out to be predominantly those already compromised and efficiently expressing the priming virus. This means that after preinfection the population of cells susceptible to a secondary infectant is largely limited to those yielding preinfectant progeny, proportion of which depends on the multiplicity of the preinfection and time interval between infections. The optimal cross protection effect may thus ensue when the proportion of cells preinfected with RSV is relatively low yet sufficient to trigger a pervasive antiviral state in bystander cells. In the case of massive infection with RSV, when most cells are simultaneously compromised by the virus, the interferon-mediated protection likely does not come into effect. The protective effect of RSV infection may increase with the yield of DVGs, which due to the lack of nonstructural proteins may both more efficiently trigger synthesis of interferons and leave STAT signaling unscathed.

The natural question is to what extent the described cross-protection mechanism reduces the incidence of coinfections or severity of the disease featured by a secondary infection *in vivo*. Answering the question is especially important, but appears complex, in the context of secondary SARS-CoV-2 infections. An early study by Ziegler *et al*. (60) showed that the receptor for SARS-CoV-2, ACE2, is a protein coded by ISG, but later Onabajo *et al*. (61) identified the transcriptionally independent truncated isoform of *ACE2, deltaACE2*, not *ACE2*, as an ISG. Since deltaACE2 may not bind SARS-CoV-2 spike protein, the latter study suggests that interferon signaling is not hijacked by this virus to enhance its proliferation. Several studies suggested beneficial roles of type I and type III interferon at the early stage of infection (62). It is thus possible that interferons secreted in response to a primary infection exhibit a protective role against a secondary infection with SARS-CoV-2 (see also (63)).

At the clinical level, one may expect that the occurrence of coinfections may result from defective work of the immune system (which aggravates the course of the disease and also imaginably cannot mediate antagonistic viral interactions). Although this effect cannot be excluded and coinfection data should be treated with caution, impact of coinfections on disease severity is unclear. Most respiratory viral coinfections, detected usually in a small proportion of patients, appear not to increase severity of clinical outcomes (64–67), with two noteworthy exceptions of a coinfection by IAV and IBV (55) and coinfections with RSV in pediatric patients (68). It should be noted that interactions of two respiratory viruses within the host can hardly be translated to their coincidence or avoidance in human populations. This is because at the population level there are two additional and antagonistic effects: on one hand, viral coincidence of contagious diseases can be seasonal (the effective contact rate is modulated by weather and behavioral patterns); on the other hand, infected and ill individuals change their behavior to avoid contact. A significant behavioral reduction of the contact rate may result from an outbreak of an epidemic of a threatening virus, decreasing the risk of propagation of other viruses, as observed during the COVID-19 pandemic in 2020 (69, 70).

In conclusion, our study demonstrates that the infection with RSV at low MOI protects alveolar epithelial cells against infection with IAV. Whereas RSV-infected cells are more vulnerable to infection with IAV, priming with RSV indirectly protects bystander non-infected cells from IAV. The cross-protection mechanism relies on induction of and paracrine stimulation with both type I and type III interferons.

## Materials and Methods

### Cell lines and cultures

The A549 and MDCK cell lines were purchased from ATTC. A549 cells were cultured in F12K basal medium supplemented with 10% fetal bovine serum (FBS) and penicillin/streptomycin antibiotic solution. MDCK cells and HEK-293T cells were maintained in DMEM medium supplemented with 10% FBS and penicillin/streptomycin antibiotic solution. All cell lines were cultured in standard conditions (37×C, 5% CO_2_) and kept in monolayers up to 90% confluency. The IFNAR1, IFNLR1, and IFNAR1–IFNLR1 knockouts based on A549 were engendered using the CRISPR lentivirus system (Abm). To obtain A549-based knockouts, HEK-293T cells were co-transfected with commercially available plasmids encoding Cas9 nuclease (Abm, cat. K002), lentivirus packaging particles (Abm, cat. LV053) and sgRNA targeted against IFNAR1 (Abm, cat. 2426411) or IFNLR1 (Abm, cat. 2487611). The double IFNAR1/IFNLR1 mutant was created based on IFNAR1 KO cells. After 2 days post-transfection, the media with lentiviral particles were collected, enriched with 8 μg/ml polybrene and filtered through 0.45 μm syringe filters. Then each individual lentiviral supernatant was used to transduce A549 wild-type cells, which were subcultured at low confluency (seeding density 5×10^4^ cells per 30 mm dish). After another 2 days, A549 cells were subjected to selection with 800 μg/ml of G418 for 10 days and seeded into a 96-well plate to obtain single cell colonies. Clones were validated by Western blot analysis of p-STAT1 and p-STAT2 in response to IFNβ or IFNλ1 (see Fig. S3). Additionally, the KO clones selected for further experiments were verified by sequencing.

### Virus amplification and isolation

Respiratory Syncytial Virus A2 strain and Influenza A virus H1N1, strain A/PR/8/34, were purchased from ATCC and amplified in HeLa or MDCK cells, respectively. Cells were seeded on a 225 cm^2^ Tissue Culture Flasks (Falcon) and cultured as described above for 2-3 days until reaching 90% confluency. On the day of infection, virus growth medium was prepared: DMEM with 2% FBS for RSV or MEM basal medium with 0.3% BSA and 1 ug/ml of TPCK-Trypsin for IAV. The dilutions of virus were prepared in an appropriate media, with target MOI around 0.01 for RSV and 0.05 for IAV. Culture media were removed, cells were washed once with PBS, and overlaid with 10 ml of inoculum. Virus was allowed to adsorb to cells for 2 hours at 37 °C with occasional stirring. Then, additional virus growth medium was added to a total volume of 40 ml per flask. Infected cells were cultured at 37 °C until the development of cytopathic effects could be observed in at least 80% of cells (typically around 3 days for IAV and 5 days for RSV). Virus-containing culture fluid was then collected and clarified by centrifugation at 3000 g, 4 °C, for 20 min. Then, virus particles were precipitated by adding 50% (w/v) PEG6000 (Sigma-Aldrich) in NT buffer (150 mM NaCl, 50 mM Tris-HCl, pH 7.5) to a final concentration of 10% and stirring gently at 4 °C for 90 min. Virus was centrifuged at 3250 g, 4 °C, for 20 min and re-centrifuged after removing supernatant to remove the remaining fluid. Pellet was suspended in 1 ml of NT buffer or in 20% sucrose in NT buffer in case of RSV, aliquoted and stored at −80 °C.

### Virus quantification

Virus concentration in collected samples was quantified using immunofluorescence protocol. HeLa or MDCK cells were seeded on microscopic cover slips and cultured upon reaching 90-100% confluency. Serial dilutions of virus samples were made in virus growth medium in a 10^−3^ to 10^−6^ range. After washing with PBS, cells were overlaid in duplicates with diluted virus, which was allowed to adhere for 2 hours with occasional stirring. Afterwards, virus-containing medium was removed, cells were overlaid with fresh virus growth medium and cultured for 16 (for IAV) or 24 hours (for RSV). Then, cells were washed with PBS and fixed with 4% formaldehyde for 20 min, RT. Cells were stained using standard immunofluorescence protocol with anti-RSV fusion glycoprotein antibody (Abcam, cat. ab43812) or anti-influenza A virus nucleoprotein antibody [C43] (Abcam, cat. ab128193). Cells containing stained viral proteins were counted using Leica SP5 confocal microscope. Virus concentration was calculated using the following formula: (avg. number of infected cells)/(dilution factor × volume containing virus added) = infectious particles/ml. For a given MOI, we observe less cells expressing viral proteins in A549 than in HeLa or MDCK lines, which are known to be permissive for, respectively, RSV and IAV infections.

### Compounds and stimulation protocols

Human interferon β1A and interferon λ1 were purchased from Thermo Fisher Scientific (cat. numbers PHC4244 and 34-8298-64, respectively) and prepared according to manufacturer’s instructions. For cell stimulation, interferons were further diluted to desired concentrations in the F12K medium supplemented with 2% FBS. Lowered FBS content was used to prevent inhibition of viral attachment and entry at the second stage of experiments.

For interferon + virus experiments, the cell culture media were exchanged for interferon-containing or control media at time = 0 and were not removed afterwards till the end of experiment. Appropriately diluted virus was added in small volumes (less than 10 μl) directly into the wells. Even distribution of virus across cell population was aided by intermittent rocking of the plate for 2 hours.

Similarly, for coinfection experiments, the F12K 2% FBS medium containing the first virus (RSV) was added to the cells at the beginning of the experiment, while the dilutions of second virus (IAV) were added directly into the wells at a later time point and distributed by rocking.

For intracellular IFNβ visualization, brefeldin A solution (BD Biosciences) was added to cells 2 hours prior to fixation.

#### Antibodies

##### Antibodies for Western blotting

Primary: anti-phospho-STAT1 (Tyr 701) (58D6) (Cell Signaling Technologies, cat. 9167, 1:1000); anti-phospho-Stat2 (Tyr 690) (D3P2P) (Cell Signaling Technologies, cat. 88410, 1:1000); anti-RIG-I (D14G6) (Cell Signaling Technologies, cat. 3743, 1:1000); anti-STAT1 (BD Biosciences, cat. 610116, 1:1000), anti-STAT2 Antibody (R&D Systems, cat. PAF-ST2, 1:1000), anti-PKR (B-10) (Santa Cruz Biotechnology, cat. sc-6282, 1:1000), anti-OAS1 Antibody (F-3) (Santa Cruz Biotechnology, cat. sc-374656, 1:1000), anti-Influenza A Virus Nucleoprotein antibody [C43] (Abcam, cat. ab128193, 1:1000), anti-Respiratory Syncytial Virus antibody [2F7] (Abcam, cat. ab43812, 1:1000); hFAB Rhodamine anti-GAPDH primary antibody (Bio-Rad, cat. 12004168, 1:10,000).

Secondary: goat anti-Rabbit IgG (H+L) Secondary Antibody, DyLight 800 4X PEG (Thermo Fisher Scientific, cat. SA5-35571, 1:10,000); goat anti-Mouse IgG (H+L) Secondary Antibody, DyLight 800 4X PEG (Thermo Fisher Scientific, cat. SA5-35521, 1:10,000); StarBright Blue 700 Goat Anti-Mouse IgG (Bio-Rad, cat. 12004159, 1:10,000), rabbit anti-goat immunoglobulins/HRP (Agilent, cat. P0449, 1:10,000).

##### Antibodies for immunostaining

Primary: anti-phospho-STAT1 Tyr 701 (58D6) (Cell Signaling Technologies, cat. 9167, 1:1000); anti-IRF-3 (D-3) (Santa Cruz Biotechnology, cat. sc-376455, 1:500), anti-Respiratory Syncytial Virus antibody (Abcam, cat. ab20745, 1:1000), anti-Influenza A virus (Abcam, cat. ab20841, 1:1000), anti-Influenza A Virus Nucleoprotein antibody [C43] (Abcam, cat. ab128193, 1:1000), anti-human interferon beta antibody (R&D Systems, cat. MAB8142).

Secondary: donkey anti-rabbit IgG (H+L), Alexa Fluor 488 conjugate (Thermo Fisher Scientific, cat. A-21206, 1:1000), donkey anti-mouse IgG (H+L), Alexa Fluor 555 conjugate (Thermo Fisher Scientific, cat. A-31570, 1:1000), donkey anti-Goat IgG (H+L) Alexa Fluor 633 conjugate (Thermo Fisher Scientific, cat. A-21082, 1:1000).

### ELISA

For measuring IFNβ production, cells were seeded on 96-well plates (Falcon) at a density of 20,000 cells per well and infected with RSV or IAV as described above. The culture medium (a total of 200 μl) from infected cells was collected at designated time points and stored at −20 °C until further analysis. IFNβ levels were estimated using the VeriKine Mouse IFNβ ELISA kit (PBL Assay Science). Standards and diluted samples in duplicates or triplicates were added to a precoated plate included in the kit and incubated for 1 h. After subsequent washing, antibody solution was prepared and added to wells for another 1 h, followed by another washing and 1 h incubation with HRP solution. Finally, TMB Substrate Solution was added to wells and developing color reaction was stopped after 15 min with the addition of Stop Solution. Optical densities of samples after resulting color development were determined using Multiskan GO plate reader (Thermo Fisher Scientific) set to 450 nm, with wavelength correction at 570 nm. IFNβ concentrations were obtained based on the standard curve (4-parameter logistic function fitted to 7 data points from standards).

### Digital PCR (dPCR)

For gene expression experiments, cells were seeded on 24-well plates at a density of 100 000 cells per well. RNA from virus-infected cells was isolated using PureLink RNA Mini Kit (Thermo Fisher Scientific), following manufacturer’s instructions: cells were harvested and vortexed in Lysis Buffer with 2-mercaptoethanol and then vortexed again with one volume of 70% ethanol. Upon transferring to the spin cartridge, cellular RNA content was bound to the column, washed with appropriate buffers and eluted, all by centrifugation at 12,000g. Eluted RNA in RNase-free water was used immediately for reverse transcription or stored for later use at −80 °C. Concentration and quality of isolated total RNA was determined by measuring UV absorbance at 260 nm and at 280 nm using Multiskan GO spectrophotometer (Thermo Fisher Scientific). Around 0.5 μg of RNA was used as a template for reverse transcription, performed using High-Capacity cDNA Reverse Transcription Kit (Thermo Fisher Scientific): diluted RNA samples were mixed 1:1 with freshly prepared Master Mix containing RT Buffer, RT Random Primers, dNTP Mix, MultiScribe Reverse Transcriptase and RNase Inhibitor. Reaction was performed in Mastercycle Gradient thermal cycler (Eppendorf) under following conditions: 10 min at 25 °C, 120 min at 37 °C, and 5 min at 85 °C.

Measurements of the viral RNA copy number was then performed using QuantStudio 3D system (Thermo Fisher Scientific) and TaqMan Gene Expression Assays Vi99990011_po and Vi99990014_po for quantification of, respectively, IAV and RSV RNA. Appropriately diluted samples were mixed with the reaction master mix and the chosen assay, loaded on the digital PCR chips in duplicates and thermocycled using ProFlex Flat Block Thermal Cycler. Subsequently, chips were analyzed with QuantStudio 3D Digital PCR Instrument. Chip quality check and final data analysis were performed using QuantStudio 3D AnalysisSuite software.

### Western blotting

Cells at the indicated time points were washed twice with PBS, lysed in a Laemmli sample buffer containing DTT and boiled at 95 °C for 10 min. Even amounts of each protein sample were separated on a 4–16% Mini-PROTEAN TGX Stain-Free Precast Gels using Mini-PROTEAN Tetra Cell electrophoresis system (Bio-Rad). Upon completion of electrophoresis, proteins were transferred to the nitrocellulose membrane using wet electrotransfer in the Mini-PROTEAN apparatus (400 mA, 50 min). Membrane was rinsed with TBST (TBS buffer containing 0.1% Tween-20, Sigma-Aldrich) and blocked for 1h with 5% BSA/TBS or 5% non-fat dry milk. Subsequently, membranes were incubated at 4 °C overnight with one of the primary antibodies diluted in a 5% BSA/TBS buffer. After thorough washing with TBST, membranes were incubated with secondary antibodies conjugated with specific fluorochrome (DyLight 800, Thermo Fisher Scientific) or horseradish peroxidase (Polyclonal Anti-Mouse/Anti-Rabbit Immunoglobulins HRP, Dako) diluted 1:5000 in 5% non-fat dry milk/TBST for 1 h, RT. Chemiluminescent reaction was developed with the Clarity Western ECL system (Bio-Rad). For GAPDH detection, hFAB Rhodamine Anti-GAPDH Primary Antibody (Bio-Rad) was used. Specific protein bands were detected using the ChemiDoc MP Imaging System (Bio-Rad).

### Immunostaining

For staining of intracellular proteins, cells were seeded on 12 mm round glass coverslips, which were previously washed in 60% ethanol/40% HCl, thoroughly rinsed with water and sterilized. After stimulation with interferon or viral infection at desired time points, cells on coverslips were washed with PBS and immediately fixed with 4% formaldehyde (20 min, RT). Cells were then washed three times with PBS and incubated for 20 min at −20 °C with 100% cold methanol. After washing with PBS, coverslips with cells were blocked and permeabilized for 1.5 hour with 5% BSA (Sigma-Aldrich) with 0.3% Triton X-100 (Sigma-Aldrich) in PBS at room temperature. After removing the blocking solution, coverslips with cells were incubated overnight at 4 °C with primary antibodies diluted in PBS containing 1% BSA and 0.3% Triton X-100. After washing cells five times with PBS, appropriate secondary antibodies conjugated with fluorescent dyes were added and incubated for 1 hour at room temperature. Subsequently, cells were washed and their nuclei were stained for 10 min with 200 ng/ml DAPI (Sigma-Aldrich). After final washing in miliQ water, coverslips with stained cells were mounted on microscope slides with a drop of Mowiol (Sigma-Aldrich). Cellular sublocalization of stained proteins was observed using Leica SP5 confocal microscope and Leica Application Suite AF software.

### Data analysis

#### Image analysis

Confocal images obtained from immunostaining were analyzed using our in-house software (https://pmbm.ippt.pan.pl/software/shuttletracker). Nuclear outlines were detected based on DAPI staining. Outlines of nuclei that were partially out of frame or mitotic were excluded from analysis; outlines of overlapping nuclei were split based on geometric convexity defects when possible. Cells were classified as either RSV-negative or RSV-positive based on the combination of the sum of intensities of pixels inside nuclear outlines (weight 5) and the perinuclear region (weight 1, decreasing sigmoidally with the distance from the nuclear contour). Cells were classified as either IAV-negative or IAV-positive based on the sum of intensities of pixels inside nuclear outlines.

#### Credible interval

Credible interval (CrI) of the Kolmogorov–Smirnov statistics (Fig. 3C, Fig. 3D, Fig. 7B, Fig. 8C, and Fig. 8D) has been estimated by resampling data points 10^5^ times. In each resampling, data points were drawn at random with probabilities proportional to the number of data points in each category (virus-positive, virus-negative) and the number of resampled data points was equal to the total number of data points in both categories.

