## Supplementary Materials (Fig. S1, Fig. S2, Fig. S3) for "RSV protects bystander cells against IAV infection by triggering secretion of type I and type III interferons"

featuring the article

by Czerkies *et al.* (*Journal of Virology*, 2022)

**Supplementary Fig. S1.** Confocal microscopy images of immunostained A549 WT cells showing activation of IRF3 (by viral RNA), phosphorylation of STAT1 (in response to interferons secreted by infected cells) and viral proteins in time points of 6, 10, 24, 48 hours post-infection with RSV at MOI of 0.01.

**Supplementary Fig. S2.** Confocal microscopy images of immunostained A549 WT cells showing activation of IRF3 (by viral RNA), phosphorylation of STAT1 (in response to interferons secreted by infected cells) and viral proteins in time points of 6, 10, 24, 48 hours post-infection with IAV at MOI of 0.01.

**Supplementary Fig. S3.** Verification of knock out cell lines derived and used in this study.

Fig. S1

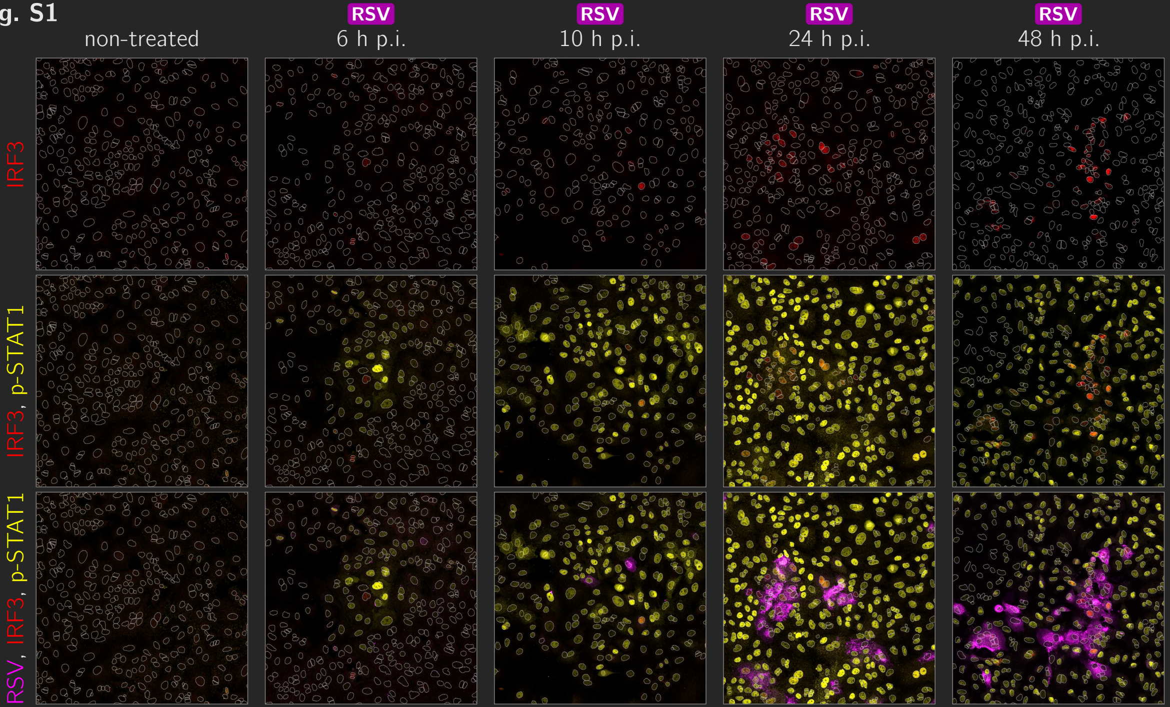

RSV
RSV
RSV
RSV
non-treated
6 h p.i.
10 h p.i.
24 h p.i.
48 h p.i.
IRF3
IRF3, p-STAT1
RSV, IRF3, p-STAT1

Fig. S2

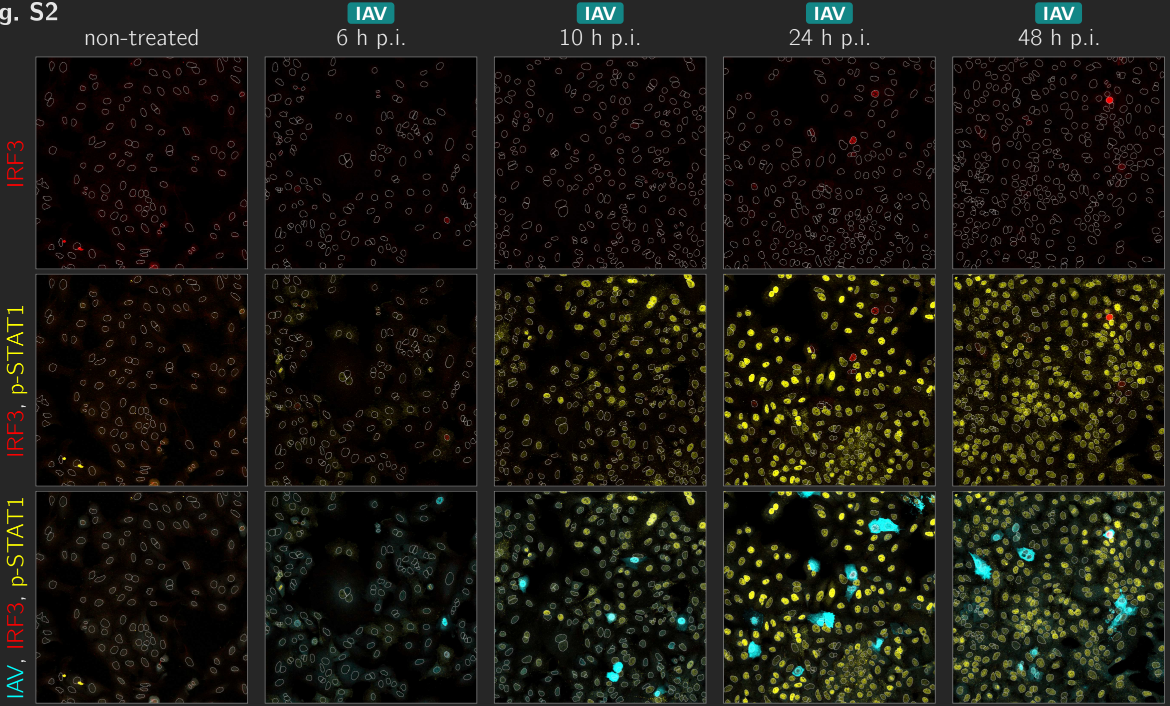

IAV
IAV
IAV
IAV
non-treated
6 h p.i.
10 h p.i.
24 h p.i.
48 h p.i.
IRF3
IRF3, p-STAT1
IAV, IRF3, p-STAT1

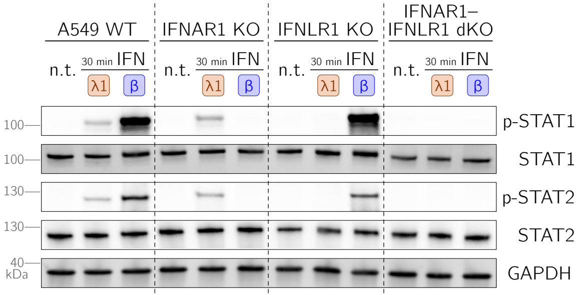


**Supplementary Fig. S3.** Verification of knock out cell lines derived and used in this study. The A549 WT, IFNAR1 KO, IFNLR1 KO, and IFNAR1-IFNLR1 double KO(dKO) cells were treated with IFNλ1 (50 ng/ml) or IFNβ (1000 U/ml) for 30 min. Non-treated cells are labeled n.t. IFN-activated STATs, p-STAT1 and p-STAT2, are STAT1 and STAT2 phosphorylated at Tyr 701 and at Tyr 690, respectively.
